# The effects of soil depth on the structure of microbial communities in agricultural soils in Iowa, USA

**DOI:** 10.1101/2020.03.31.018416

**Authors:** Jingjie Hao, Yen Ning Chai, Raziel A. Ordóñez, Emily E. Wright, Sotirios Archontoulis, Daniel P. Schachtman

## Abstract

The determination of how microbial community structure changes within the soil profile, will be beneficial to understanding the long-term health of agricultural soil ecosystems and will provide a first step towards elucidating how deep soil microbial communities contribute to carbon sequestration. This study aimed to investigate the differences in the microbial community abundance, composition and diversity throughout from the surface layers down to deep soils in corn and soybean fields in Iowa, USA. We used 16S rRNA amplicon sequencing of soil samples to characterize the change in microbial community structure. Our results revealed decreased richness and diversity in bacterial community structure with increasing soil depth. We also observed distinct distribution patterns of bacterial community composition along soil profiles. Soil and root data at different depths enabled us to demonstrate that the soil organic matter, soil bulk density and plant water availability were all significant factors in explaining the variation in soil microbial community composition. Our findings provide valuable insights in the changes in microbial community structure to depths of 180 cm in one of the most productive agricultural regions in the world. This knowledge will be important for future management and productivity of agroecosystems in the face of increasing demand for food and climate change.

## Introduction

Microbial communities play pivotal roles in the ecosystem, plant and animal health, food safety and climate change factors [1–3]. Soils are one of the most diverse ecosystems on Earth containing microscopic bacteria and fungi, microfauna (nematodes and protozoans), mesofauna and macrofauna [4]. Soil microbiomes are a foundational feature of the agricultural ecosystems and host various biogeochemical processes, such as soil nutrient cycling and organic matter decomposition. Soils also provide plant roots with the nutrients required for growth and productivity, store carbon, and emit greenhouse gases that contribute to climate change [5–8]. Despite the extensive presence of soil microbes throughout the soil profiles, current understanding of the diversity and composition of soil microbial communities is mainly restricted to the top surface of soils (∼ 0 - 25 cm), where there are higher proportions of soil nutrients, organic matter and a higher diversity of microorganisms than that in the subsurface layers [9,10]. However, in rainfed production systems the entire soil profile is important for crop yield. This is because the topsoil can dry out very fast during summer months if no rain occurs. This can limit the ability of roots to absorb water and nutrients. Therefore, extending our microbial knowledge to deeper depths is important.

The composition of soil microorganisms is influenced by a variety of edaphic factors, such as habitat type, soil pH, texture, moisture, mineral nutrient content and organic matter [11–16]. Previous studies have demonstrated remarkable changes in microbial community composition with soil depth across different environments [10,17–20], such as consistent decreases in microbial abundance and diversity with increasing depth [21]. The differences of microbial community composition between surface soils and subsurface soils have been attributed to the drastic differences in soil nutrients, extracellular enzyme activities, soil organic carbon, and microbial biomass [22,23]. However, most of these studies focused on the microbial diversity variation between the surface and subsurface soils at a depth range of 0 - 100 cm in natural soils [24], in very specific environments, such as Alaskan soil cores or in a paddy soils [11,19]. The subsurface soil microbial communities are also very important to characterize because they have greater impact on soil forming processes than surface soils [25]. Microbial communities in subsurface soils may also play important roles in soil carbon sequestration because of the enrichment of organic carbon stored in the subsurface profiles [26,27]. Therefore, exploring the characteristics of subsurface soil microbial communities throughout the soil profiles will enable us to better understand multiple soil processes, some of which are significant components in carbon cycling that can mitigate climate change [28].

In addition to soil depth, plants are another key factor that influence soil microbial activities in various ecosystems [29,30]. Since plant growth is tightly interrelated with edaphic factors, the plant-soil interactions at different soil depths play a role in affecting the abundance and composition of soil microbial communities [31]. Although most studies focused on the nutrient rich topsoil, the roots of agricultural crops can grow as deep as 200 cm [32]. For example, the average maximum rooting depth of corn and soybean grown in the midwestern USA is 150 cm [33]. Soil depth has been identified as a critical edaphic factor that can shape soil microbial community composition in arable soil [34]. Changes in different bacterial key taxa that utilize plant-derived carbon in the rhizosphere of wheat at different depths in an arable soil have been reported [35]. However, our understanding about the vertical distributions of soil microbiomes in agricultural soils is very limited. Investigating the soil bacterial biogeography along the field crop rooting systems, especially in deeper soil profiles, will provide insights into distinct and potentially important processes involved in agricultural soil ecosystems, which in turn would also assist developing better crop management strategies for specific crops, soil type and climate factors.

Crop production in agriculture fields requires effective agronomic management practices. Understanding the soil microbial properties with respect to changes in soil depth could be potentially beneficial to the long-term health of agricultural soils. This study was carried out to investigate the effects of depth on the microbial community abundance, composition and diversity in soils collected from corn and soybean fields in Iowa, which is located in one of the world’s most productive agricultural regions [36]. We used a dataset of 16S rRNA amplicons of soil DNA that was sequenced on an Illumina MiSeq platform. Our objective was to investigate the differences in the composition of soil microbial communities throughout the soil profiles in these agricultural fields.

## Materials and methods

### Field sites and soil sample information

Soil samples were collected from Ames, Kelley, and Kanawha located in Des Moines lobe of Iowa, USA. Soybean (*Glycine max*) and corn (*Zea mays*) were planted in Ames and Kelley, while corn was the only crop in Kanawha. The Kelley site had subsurface tile drainage installed at 1.1 m below the surface, which meant that the 0 - 1 m soil profile rarely is saturated with water. In contrast, Ames and Kanawha that shared the same soils with Kelley (Nicollet soil series), had no tile drainage, so that the 0 - 1 m profile was saturated with water for longer periods. Recent experimental and modeling studies carried out in these fields showed that the depth of the water table and the hydrology of the field dictates the corn and soybean root distribution. The Kelley site has been under no-tillage management since 2009, the other two sites were tilled every autumn. Detailed description of soil properties, and management practices is provided by Archontoulis *et al*., [37], Nichols *et al*., [33], and Ordóñez *et al*., (under review). Soil cores were collected on the plant row and between two rows (36 cm apart between sampling points) during mid grain filling period (a period when root mass is maximum) from 0 - 210 cm in corn and soybean fields using Giddings probe (6.2 cm diameter). Soil samples were always kept under fresh condition at all times to maintain sample quality. Before submitting samples for washing and determination of root properties, each sample was well mixed, and a sub-sample was extracted for microbial analysis. Soil samples from the 180 - 210 cm depth layer yielded very few microbial DNAs and were excluded from our dataset. The deep cores were sectioned into seven depth intervals (0 - 15, 15 - 30, 30 - 60, 60 - 90, 90 - 120, 120 - 150 and 150 - 180 cm).

### DNA extractions, 16S rRNA gene amplification and sequencing

DNA was extracted from soil samples using the PowerSoil-htp 96 Well Soil DNA Isolation Kit (MoBio, Carlsbad CA). The V4 region of the 16S rRNA gene was amplified by PCR using a dual-index sequencing strategy [38] with AccuPrime™ *Pfx* DNA Polymerase (Invitrogen, Carlsbad CA). A dual-index primer system was used and consists of the Illumina adapter, an 8 - nucleotide index sequence, a 10 - nucleotide pad sequence, a 2 - nucleotide linker sequence, and the 16S V4 - primer [38]. Amplification reactions were checked by running PCR products on 1% agarose gel to ensure success of PCR. The PCR reactions were purified and normalized using the SequalPrep™ Normalization Plate Kit (Invitrogen, Carlsbad CA). The concentration of PCR products was then measured using the QuantiFluor™ dsDNA System (Promega, Madison, WI) and used to pool equimolar amounts of PCR products together. Pooled samples were concentrated using a SpeedVac, and fragments within a size range of 200 - 700 bp were size-selected using the SPRIselect beads (Beckman Coulter, Brea, CA). In the amplicon library, a blank DNA extraction control was used as a negative control. Genomic DNA from Microbial Mock Community B (Even, Low Concentration), v5.1L for 16S rRNA Gene Sequencing (BEI Resources, Manassas, VA) was used as a positive control. Sequencing libraries were quantified, and quality checked using a High Sensitivity DNA kit on an Agilent 2100 Bioanalyzer (Agilent Technologies, Santa Clara, CA). Sequencing was performed on the Illumina MiSeq platform using the MiSeq Reagent Kit v3 (600 cycles, Illumina) with a spiking of 20% PhiX control library (Illumina, San Diego, CA).

### Sequence processing and data analysis

The raw paired-end sequencing reads were processed using USEARCH (version 10.0.240) and QIIME (Quantitative Insights into Microbial Ecology, version 1.9.1) [39]. Briefly, sequence reads were de-multiplexed and high-quality merged reads were clustered with simultaneous chimera removal using UNOISE implemented in USEARCH into amplicon sequence variant (ASVs) based on 100% sequence similarity. OTUs were classified using the Ribosomal Database Project (RDP) classifier [40] against the GreenGenes 16S database [17]. Chloroplasts and mitochondria sequences were identified and removed from the data. Low abundance OTUs (< 2 total counts) were discarded. All samples were rarefied to 5,024 sequence reads per sample and samples having sequences fewer than 5,024 were removed. The microbial alpha diversity was evaluated by calculating observed OTUs and Shannon diversity index, and the microbial beta diversity was assessed by calculating the Bray-Curtis dissimilarity between samples. Canonical analysis of principal coordinates (CAP) analyses were conducted using the ‘capscale’ function in vegan (v2.5.3) R package [41]. Data visualization was performed using the ggplot2 (v2.2.1) [42]. The taxa shift along with different depths were presented in bar plots based on the percent relative abundances of the top 20 most abundant microbes at the phylum level.

### CCA analysis

Canonical correspondence analysis (CCA) was conducted to explore the relationship between the bacterial community composition and soil chemical properties (soil physicochemical variables listed in Supplementary Table S1) which were partially reported in an earlier study [33] using the ‘cca’ function in R [41]. Soil properties that led to statistically significant change in microbial community composition were selected to build the CCA model using ‘ordistep’ function with 999 permutations. Statistical significance of each soil chemical property and CCA axes were determined using Monte Carlo permutation test with 999 permutations.

### Statistical analysis

Differences in microbial alpha diversity were determined using one-way ANOVA followed by a pairwise Tukey’s HSD tests with JMP Pro 13 software (SAS Institute Inc., Cary, NC). Permutational multivariate analysis of variance (PERMANOVA) analysis was performed to assess the effects of soil depth, sites, and crop type on bacterial community data using the ‘adonis’ function in vegan R package [41]. Pairwise comparisons of Bray-Curtis dissimilarities between corn and soybean soil across soil depth were conducted using two-sided Student’s two-sample t-test. *P* < 0.05 was considered as statistically significant.

## Results

### Decreased richness and diversity in bacterial community along soil depth gradient

Microbial richness as determined by observed OTUs was highest in the surface soil and decreased as soil depth increased. The richness of bacterial community was similar between 15 - 30 cm and 30 - 60 cm, 30 - 60 cm and 60 - 90 cm, 90 - 120 cm and 120 - 150 cm, 120 - 150 cm and 150 - 180 cm (**Fig 1A**). The diversity of the bacteria community resembled each other between 0 - 90 cm, but a decreasing trend in diversity was observed between 90 - 180 cm (**Fig 1B**).

**Fig 1.**
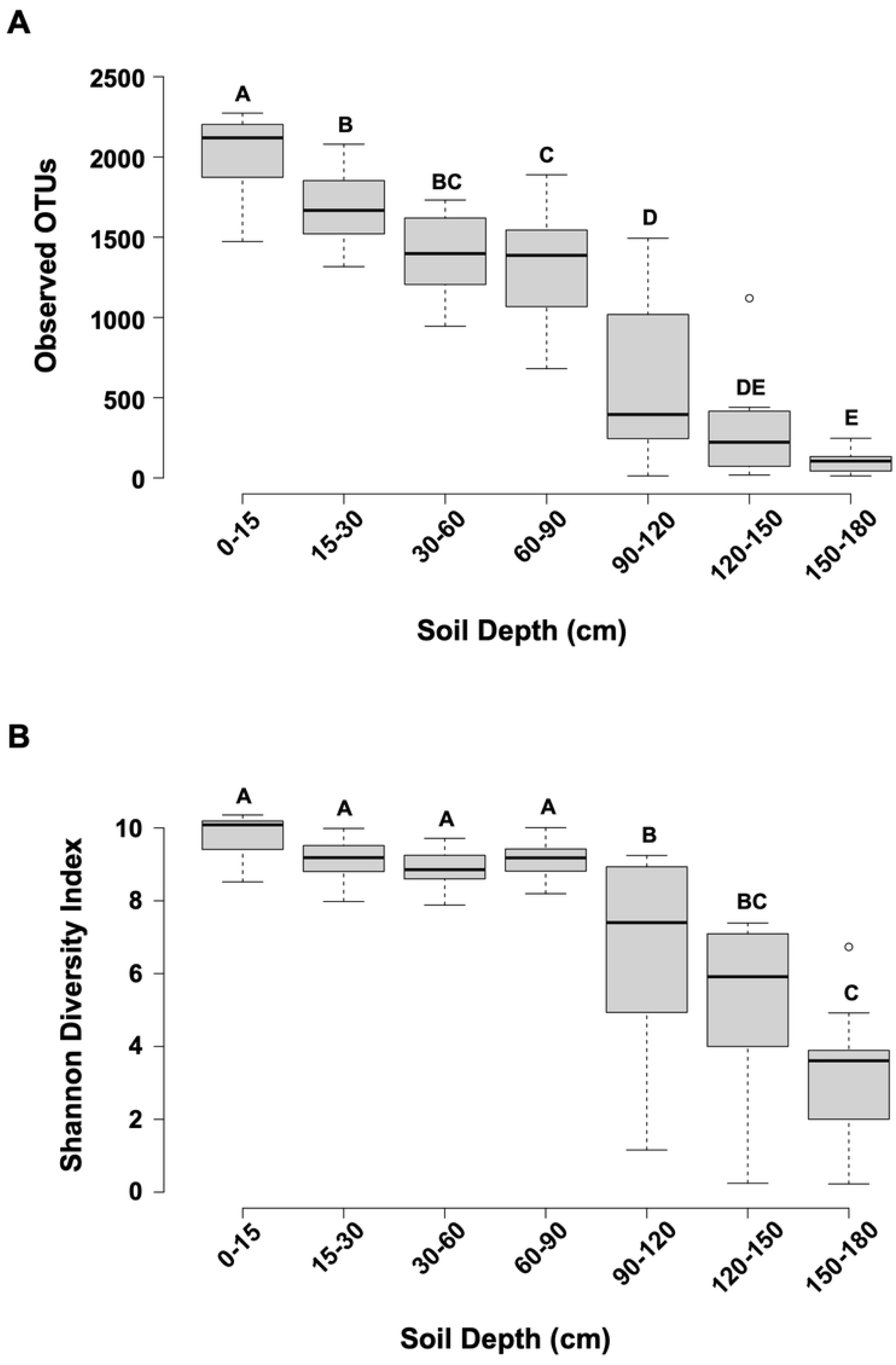
Changes in alpha diversity levels with depth. (A) Averaged number of observed OTUs at different soil depth. (B) Shannon diversity at different soil depth. Differences in alpha diversity were compared using one-way ANOVA followed by Tukey’s HSD tests. *P* < 0.05 was considered as statistically significant. Different letters above the bars denotes significant differences between soil depths. Line in the box represents median. The top and bottom of box represent the first and the third quartile, respectively. Whiskers indicate data’s range.

### Soil depth shifts the microbial community composition

Canonical analysis of principal coordinates (CAP) was performed to evaluate how each variable in the dataset, including soil depth, sampling site and crop type, contributed to the variation in microbial community composition. Soil bacterial community composition shifted significantly with soil depth (*p* < 0.001, 31.04% variation explained), sampling sites (*p* < 0.001, 4.01% variation explained), and crop types (*p* < 0.05, 1.23% variation explained) (**S1 Fig**). In addition, there was a significant interaction between depth and site (*p* < 0.05, 10.59% variation explained) (**S1 Fig**). To assess the influence of soil depth alone on soil bacterial community composition, CAP analysis was performed based on a Bray-Curtis dissimilarity factoring out the effect of site and crop type. Those results showed that the bacterial community composition was significantly different among samples at different soil depths (*p* < 0.001) (**Fig 2**). The CAP analysis was also conducted using both weighted UniFrac (WUF) and unweighted UniFrac (UUF) distance metrics and similar results were obtained (data not shown).

**Fig 2.**
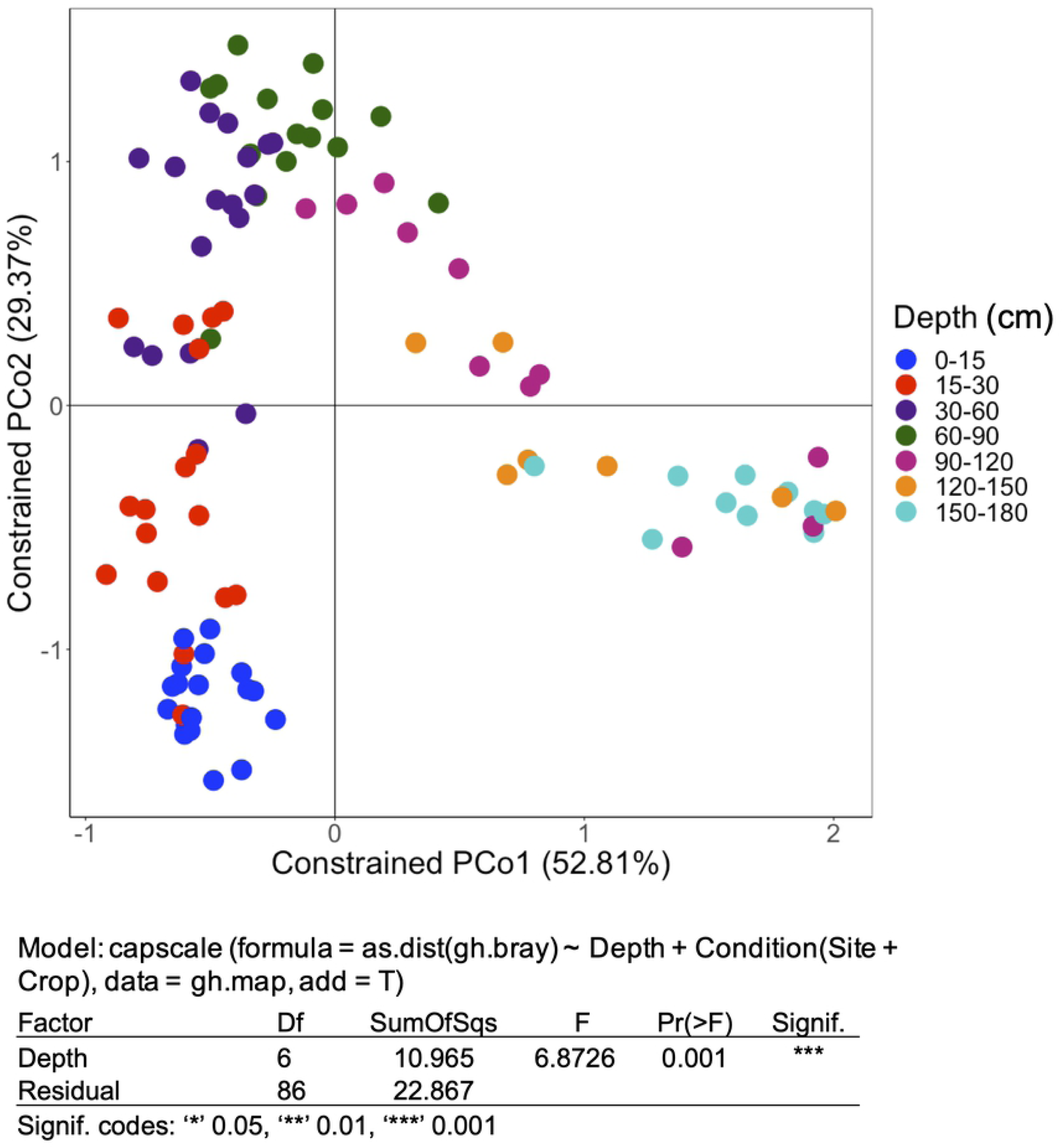
Beta diversity showing changes in microbial community composition with depth. Canonical analysis of principal coordinates (CAP) using Bray-Curtis dissimilarity for all samples. The Bray-Curtis dissimilarity matrix was generated using QIIME. CAP analysis was conducted by constraining soil depth using the ‘capscale’ function in vegan R package. PERMANOVA statistical analysis was performed to determine whether the differences between soil depths was significant. Each color indicates a different soil depth as shown in the legend.

**S1 Fig. Beta diversity showing changes in microbial community composition with depth, site and crop type**. Canonical analysis of principal coordinates (CAP) using Bray-Curtis dissimilarity for all samples. The Bray-Curtis dissimilarity matrix was generated using QIIME. CAP analysis was conducted by constraining soil depth, crop type and sampling site using the ‘capscale’ function in vegan R package. PERMANOVA statistical analysis was performed to determine whether the shifts in bacterial community due to soil depth, crop type, sampling site, and their interactions were significant. Each color indicates different soil depth as shown in the legend.

### Effect of crop type and sampling location on the microbial community composition across soil depth

To assess if crop type or sampling location influenced microbial community composition at different soil depths, we used a pairwise comparison of Bray-Curtis dissimilarities between the soils from corn and soybean fields (**Fig 3A**) or among the three sampling locations including Ames, Kelley and Kanawha (**Fig 3B**) along soil depth gradient. Soil bacteria community composition was significantly different between the two crop types at the top three surface soil layers including 0 - 15 cm (*P* ≤ 0.01), 15 - 30 cm (*P* ≤ 0.001) and 30 - 60 cm (*P* ≤ 0.001), and resembled each other at soil depths deeper than 60 cm (**Fig 3A**). More variation of soil bacterial community composition was observed across the three sampling locations to a soil depth of 90 cm, while no significant variation was observed in deeper soil layers. The soil bacterial community of Kanawha was consistently different from Ames at soil depths of 0 - 90 cm, and different from Kelley at depths 15 - 90 cm (Fig 3B). Ames and Kelley had distinct bacterial communities only in the topsoil layers (0 - 30 cm). These results indicated that both crop type and sampling location had significant effect on the bacterial community composition in the upper profile of the soil.

**Fig 3.**
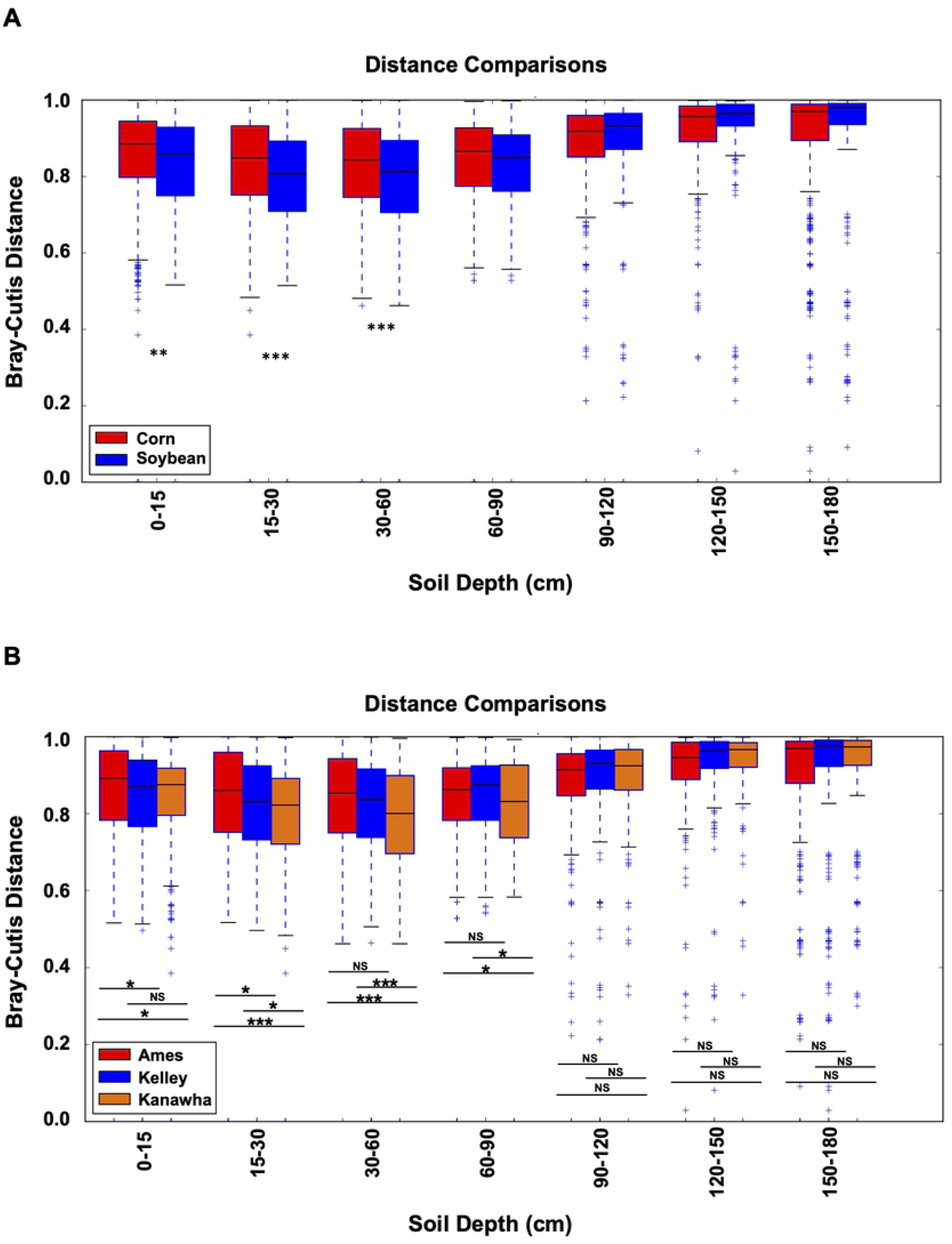
Distribution of pairwise Bray-Curtis dissimilarities between crop type along soil depth gradient. (A) Bray-Curtis distances between soils from corn and soybean fields and (B) between soils from each of two locations along a soil depth gradient were computed using ‘make_distance_comparison_plots.py’ function in QIIME 1. Significance tests were performed using two-sided Student’s two sample t-test. Asterisks indicate significant differences (**, 0.01; ***, 0.001). NS, not significant.

### Soil properties correlated with microbial community composition

The canonical correspondence analysis (CCA) showed that soil depth was the most dominant (*P* ≤ 0.001) factor shaping the microbial community composition, explaining 15.4% of the variation in microbial communities (**Fig 4**). Sampling site was also significant in this analysis (*P* ≤ 0.001). Among the soil properties analyzed, soil organic matter (*P* ≤ 0.001), bulk density (*P* ≤ 0.05), and the length of time the subsurface soil was inundated by water (*P* ≤ 0.001) were significant in explaining the variation in soil microbial community composition. The soil organic matter was responsible for 2.1 % of the total variation in bacterial community and length of time the subsurface soil was inundated by water explained 1.5% (**Table 1**). Other variables including root biomass, root length and plant water availability were not statistically significant (*P* > 0.05) in influencing the soil microbial communities.

**Table 1.** Statistical analysis for CCA. Soil properties led to statistically significant changes in microbial community composition, including ‘days saturated’, ‘organic matter’ and ‘bulk density’ were selected to build the CCA model. *p* < 0.05 was considered as significantly different.

|  | Df | ChiSquare | F | Pr(>F) |  |
| --- | --- | --- | --- | --- | --- |
| Depth | 6 | 1.9562 | 3.8015 | 0.001 | *** |
| Site | 2 | 0.3876 | 2.2598 | 0.001 | *** |
| Days Saturated | 1 | 0.2003 | 2.3349 | 0.001 | *** |
| Organic Matter | 1 | 0.1558 | 1.8165 | 0.001 | *** |
| Bulk Density | 1 | 0.1425 | 1.6610 | 0.002 | ** |
| Crop | 1 | 0.1033 | 1.2043 | 0.086 |  |
| Root Length | 1 | 0.1249 | 1.4558 | 0.075 |  |
| Root Biomass | 1 | 0.0819 | 0.9545 | 0.451 |  |
| Plant Available Water | 1 | 0.1105 | 1.2887 | 0.106 |  |
|  | 111 |  |  |  |  |
| Residual |  | 9.4342 |  |  |  |
| Signif. codes: 0 '***' 0.001 '**' 0.01 '*' 0.05 |  |  |  |  |  |

**Fig 4.**
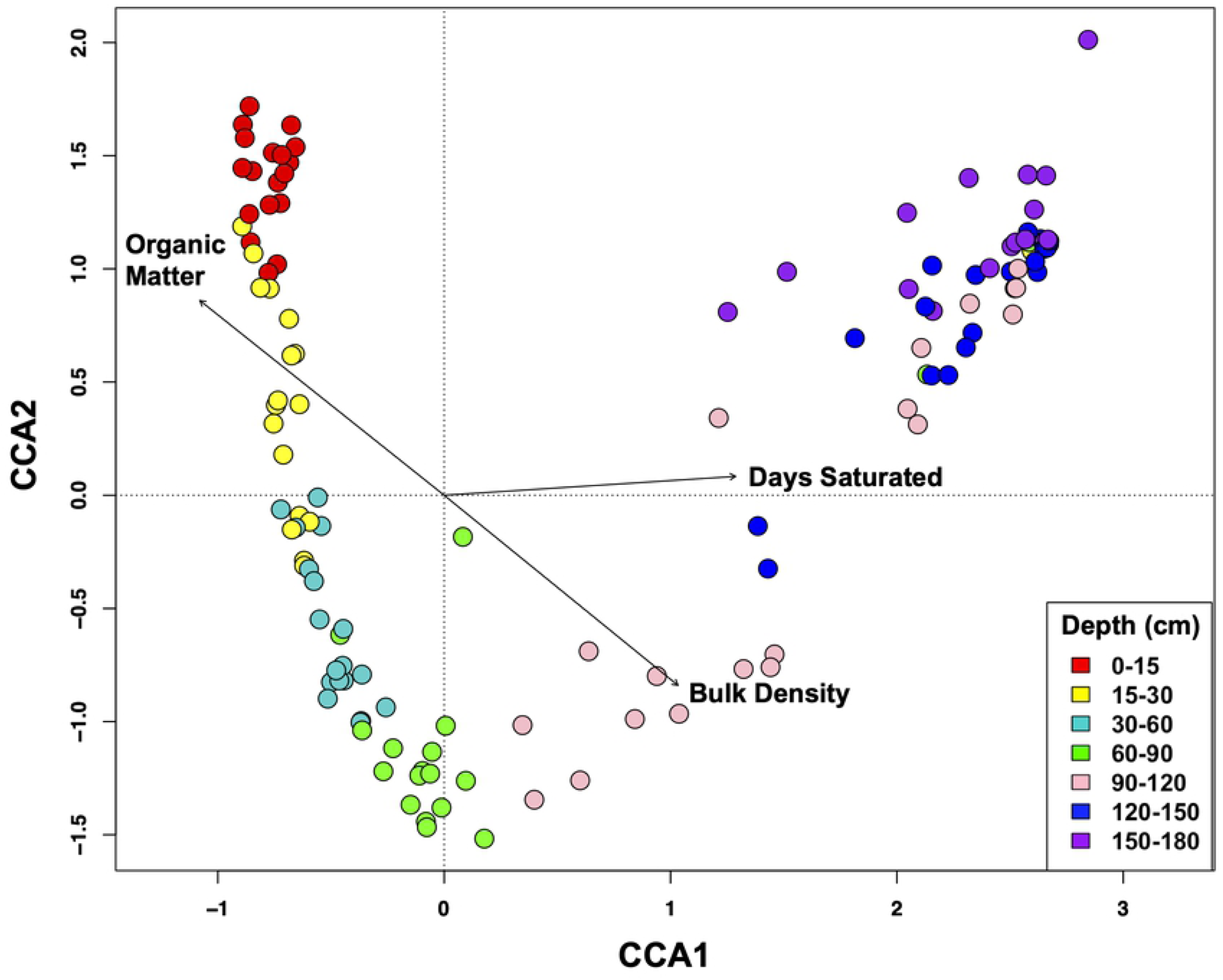
Canonical Correspondence Analysis (CCA) to examine the additional factors influencing soil microbial community composition. CCA1 is the constrained ordination of the data with 69.01% (*p* < 0.001) of the variation and CCA2 with 45.00% (*p* < 0.001) of the total variation of each axis. The significance for each soil property is presented in Table 1.

### Changes in specific bacterial taxa along soil depth

A total of 31,230 OTUs were retrieved and assigned to 53 phyla, 138 classes, 208 orders, 238 families and 306 genera in all 126 soil samples. The dominant bacterial phyla of the microbial communities include *Proteobacteria, Actinobacteria, Acidobacteria, Chloroflexi, Planctomycetes, Verrucomicrobia, Crenarchaeota, Nitrospirae*, which accounted for more than 89.7% of bacterial sequence reads (**Fig 5**). The relative abundance of *Acidobacteria* gradually declined with depth. The phylum *Actinobacteria* increased in relative abundance with depth in the top 0 - 60 cm, peaking in the 30 - 60 cm depth soil layer, and decreasing gradually with depth in the deeper horizons (60 - 180 cm). The phylum *Verrucomicrobia* and *Crenarchaeota* were more abundant in the surface soil layers (0 - 60 cm) than that in the deeper regions (60-180 cm), although they accounted for a relatively small proportion of the bacterial community in all soil layers. The relative abundances of *Planctomycetes* did not show clear shifts across different depths, except in the very deep region from 150 - 180 cm, where relative abundance noticeably decreased (**Fig 5**, Supplementary Table 2). Taken together, these results show the changes in specific bacterial phylum that contribute to the shift in bacterial community composition due to soil depth.

**Fig 5.**
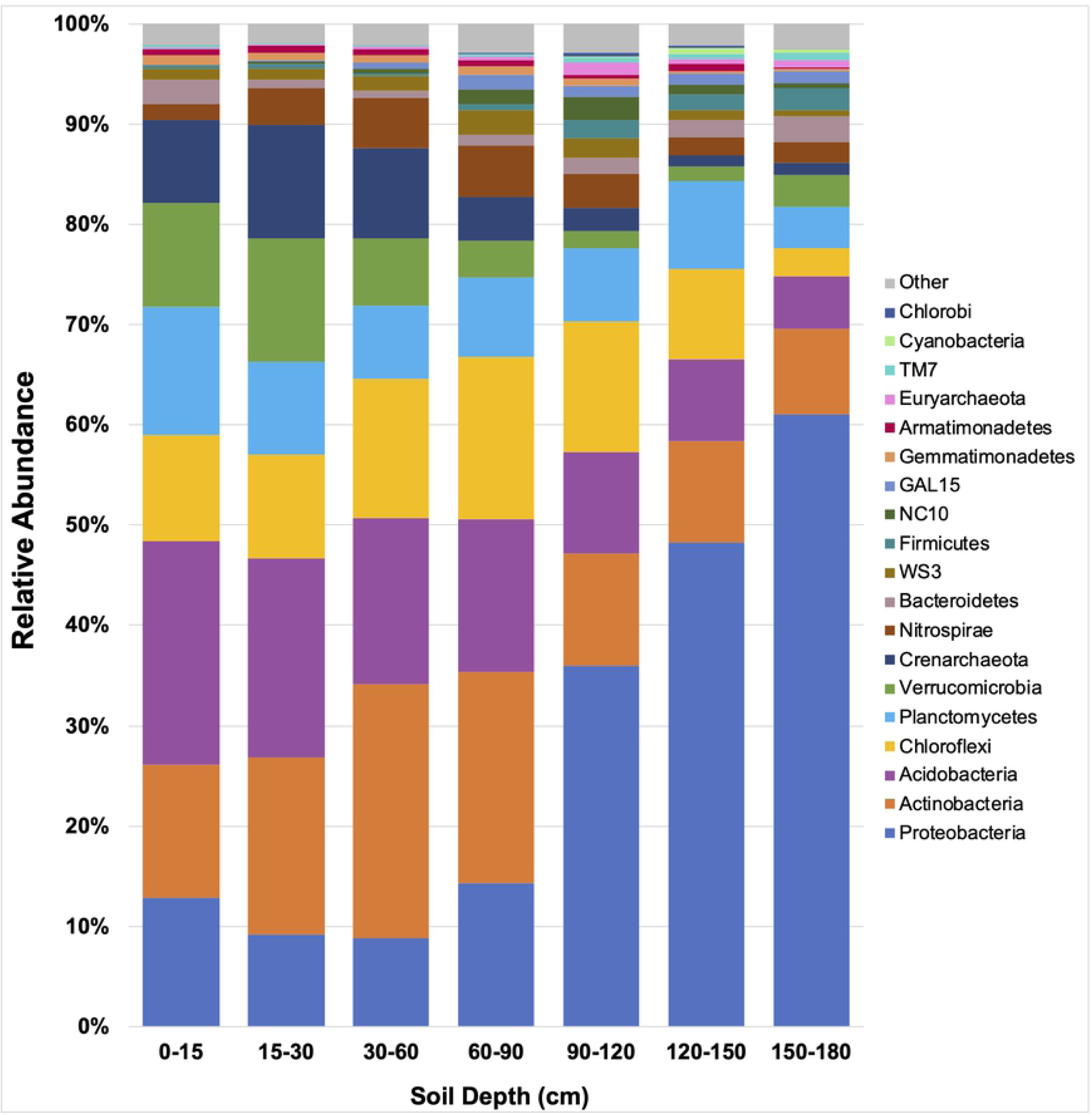
Relative abundance of the dominant microbial communities in all samples separated by soil depth. Bar plots of phylum-level relative abundance of the top twenty most abundant taxa.

## Discussion

This study demonstrated that soil profiles, representing changes of various edaphic factors with depth (i.e. soil carbon, bulk density, water-holding capacity), had significant effects on the composition of microbial communities in agricultural soils. Our results are consistent with previous findings that soil depth is a fundamental environmental factor in structuring soil microbial communities [43]. Decreased richness and alpha diversity in bacterial community along soil depth gradient were also observed in the present study. Furthermore, our study used a dataset obtained from corn and soybean cropping systems [33], providing new evidence that the effect of soil depth on structuring subsoil microbial communities is also very important in the agroecosystems.

The association of plant roots and soil microbial communities has great influence on plant growth, health and stress tolerance [44]. It also plays important roles in biogeochemical processes of the Earth such as carbon sequestration, water cycling, as well as emission of greenhouse gases [2,45]. In agroecosystems, it is important to characterize the spatial distribution of soil microbiota along soil profiles as the vertical distribution of crop roots system has a significant effect on soil microbial communities [46]. Emerging evidence has indicated that “digging deeper” to assess the impact of deep roots on soil microbial communities is essential to enhance our understandings of community ecology and ecosystem functioning [24]. Soil is the major reservoir of organic carbon on earth, and therefore the components affecting carbon storage and release from soils are critical for our full understanding of the role of different soil types in climate change [47]. The microbial communities play important roles in the turnover and transport of greenhouse gases, which in turn contributes to mitigation of climate change in terrestrial and aquatic ecosystems [48,49]. Although the carbon content is relatively less in subsoil horizons, it still accounts for more than 50% of the total soil carbon, therefore it is important in the global carbon cycle [50]. It has been reported before that microbes in deeper soils have a crucial influence on soil formation and soil carbon sequestration [27]. Our high-resolution root and soil data at different depths enables us to demonstrate that the soil organic matter, soil bulk density and plant water availability are both significant factors in explaining the variation in soil microbial community composition.

Site location was the second significant factor that impacted soil bacterial communities, after soil depth. However, the influence of sites on soil bacterial communities was only significant at topsoil from 0 to 90 cm, and not at deeper soil layers. By comparing the Bray-Curtis dissimilarity at different soil depths, we found that at the Kanawha site soil bacterial composition differed in the shallower soil depths compared to that of the Ames and Kelley sites. The soil bacterial communities between Kanawha and Ames sites differed significantly down to a soil depth of 90 cm, while the Kanawha and Kelley sites exhibited different bacterial communities from 30 to 90 cm. This could be attributed to the fact that Kanawha is geographically (northern) farther away from Ames and Kelley, while Ames and Kelley are located closer together (about 5 miles distance). Despite sharing similar soil carbon and nitrogen among these three sites, the weather was very different between Kanawha and the other two sites. In Kanawha, the amount of precipitation during the growing season was greater while the average temperature was lower than the Ames and Kelley sites, which might explain why the Kanawha site had a more distinct soil microbial community in the topsoil than the other two sites since topsoil is more susceptible to climate differences compared to deeper soil. Two studies suggested that different tillage management and soil hydrology can drive the differentiation of the soil microbiome in the topsoil [51,52]. Our analysis also found significant association between the length of time soil was saturated with water and plant water availability differences in bacterial community composition among these three sites (Fig 4; Table 1).

In addition to depth and site, crop type contributed significantly to the variation in bacterial communities only at the topsoil layers. These results are consistent with a previous study investigating the effect of biofuel crops on soil microbial communities at different soil depths from 0 - 100 cm [53]. In the present study, soybean and corn root biomass from 0 - 90 cm was twelve-fold greater than at 90 - 180 cm, thus richer in plant root exudates and therefore greater organic matter deposition is expected in the topsoil than in the deeper soil layer. Plant root exudates are metabolites that shape the microbial communities and differ between plant species [54,55] and within species [56]. Therefore, the crop type effect on soil bacterial communities might be due to the fact that soybean and corn had exudates that differ in composition and therefore influenced the structure of the bacterial communities. The effect of crop type on soil bacterial communities was not detected at deeper soil depths potentially because the root mass was much less in deeper soils. In alignment with that, no significant difference was found in root mass below 90 cm soil profile when comparing both crops at each site (*P* > 0.22). Intensifying substrate and oxygen limitations in deeper soil layers also impose a selective pressure on the bacterial communities, reducing their diversity and richness compared to the top soil layers [57].

Previous studies show evidence that bacterial phyla change in relative abundance with soil depth [10], although the patterns of change of individual bacterial taxa are highly variable among different studies. For example, the relative abundance of *Actinobacteria* and *Acidobacteria* in one study fluctuates throughout the soil profile [58]. In our present study, the relative abundance of *Acidobacteria* consistently declined with increasing of soil depth, while *Actinobacteria* peaked in relative abundance in the 30 - 60 cm layer. Another study focusing on the influence of long-term fertilization on changes in deep soil microbial communities showed that *Proteobacteria* was the dominant phylum in deep soil, e.g. about 84% in the 2 - 3 m layer of the soil profile [59]. We also observed that *Proteobacteria* was the most dominant phylum in subsoils, accounting for about 35% - 62% of relative abundance in deep layers of soil profile (90 - 180 cm). The reason for the abundance of *Proteobacteria* with the increasing of depth could not be elucidated in this study. Together, these inconsistent findings among different studies suggest that changes in microbial community composition with soil depth depend on the geographic locations and specific features of the soil profiles being studied.

The development of a better understanding of changes and factors that influence plant-microbe-soil interactions through the soil profile in the agroecosystems will enable us to better deploy plant and microbial solutions to improve crop yields, enhance carbon storage and allocation, and greenhouse gas mitigation, in response to population growth, increased demand for food, as well as climate change.

## Acknowledgements

Thanks to Lucas Dantes Lopes and Ellen Marsh for comments on the manuscript, to Stephanie Futrell for excellent technical assistance, and to the University of Nebraska Medical Center Genomics Core for sequencing. This work was supported in part by a grant from the National Science Foundation EPSCOR Center for Root and Rhizobiome Innovation Award OIA-1557417, Foundation for Food and Agricultural Research (Grant #534264) and USDA Hatch project (IOW10480).

